# Identification of conserved and tissue-restricted transcriptional profiles for lipid associated macrophages (LAMs)

**DOI:** 10.1101/2024.09.24.614807

**Authors:** Yingzheng Xu, Hannah Hillman, Michael Chang, Stoyan Ivanov, Jesse W. Williams

## Abstract

Macrophages are essential immune cells present in all tissues, and are vital for maintaining tissue homeostasis, immune surveillance, and immune responses. Considerable efforts have identified shared and tissue-specific gene programs for macrophages across organs during homeostasis. This information has dramatically enhanced our understanding of tissue-restricted macrophage programming and function. However, few studies have addressed the overlapping and tissue-specific responses of macrophage subsets following inflammatory responses. One subset of macrophages that has been observed across several studies, lipid-associated macrophages (LAMs), have gained interest due to their unique role in lipid metabolism and potential as a therapeutic target. LAMs have been associated with regulating disease outcomes in metabolically related disorders including atherosclerosis, obesity, and nonalcoholic fatty liver disease (NAFLD). In this study, we utilized single-cell RNA sequencing (scRNAseq) data to profile LAMs across multiple tissues and sterile inflammatory conditions in mice and humans. Integration of data from various disease models revealed that LAMs share a set of conserved transcriptional profiles, including *Trem2* and *Lpl*, but also identified key sets of tissue-specific LAM gene programs. Importantly, the shared LAM markers were highly conserved with human LAM populations that also emerge in chronic inflammatory settings. Overall, this analysis provides a detailed transcriptional landscape of tissue-restricted and shared LAM gene programs and offers insights into their roles in metabolic and chronic inflammatory diseases. These data may help instruct appropriate targets for broad or tissue-restricted therapeutic interventions to modulate LAM populations in disease.

## Introduction

Macrophages are phagocytic immune cells, and are present in all tissues where they play diverse roles in tissue homeostasis, surveillance, and clearance of cellular debris^1^. While macrophages are recognized to have conserved transcriptional profiles, tissue-specific gene programs have been described in tissue resident macrophages across tissues^2,3^. Since foundational studies that describe unique tissue-specific transcriptional programs, several groups have investigated the functional regulation of these genes and their contribution to promoting organ-specific functions^4–6^. As such, several recent studies have further resolved conserved macrophage profiles across tissues in the steady state^7–9^. Beyond these studies in homeostatic settings, it was suggested that conserved gene programs emerged following tissue injury. However, limited studies have investigated the conserved or tissue specific programming of the macrophage subsets that emerge during tissue injury.

One macrophage subset recently recognized to expand following sterile tissue injury in several settings are lipid associated macrophages (LAMs). LAMs are a unique subset of macrophages that play a pivotal role in lipid metabolism and tissue homeostasis^10–12^. These macrophages are increasingly appreciated in recent years due to their influence in various diseases of metabolic dysregulation, including atherosclerosis, obesity, nonalcoholic fatty liver disease (NAFLD), and cancer^13–19^. Understanding the biological functions of LAMs is essential to develop therapeutic strategies for these conditions.

Multiple studies have shown that LAMs can originate from tissue resident macrophages or circulating monocytes that enter tissue in response to inflammatory cues, and differentiate into terminal cells with specific lipid-handling abilities, including sensing, engulfing, efflux, and breaking down of lipids^11,20–25^. These cells also exhibit unique lipid associated markers transcriptomic profiles that distinguish them from other macrophage subsets. For instance, several studies have identified that LAMs are characterized by the expression of markers such as Triggering Receptor Expressed on Myeloid cells 2 (Trem2) and Lipoprotein Lipase (Lpl), emphasizing their role in lipid sensing and metabolism. Specifically, LAMs help take up and break down excessive lipid to prevent lipid accumulation that may lead to metabolic disorders. This homeostatic function contributes to metabolic and immune stability of tissues and ensures healthy energy storage and usage^26–29^.

LAMs are documented in multiple tissue sites, including adipose tissue, liver, adrenal gland, arterial plaques, and even the tumor microenvironment. In the adrenal gland, LAMs closely interact with lipid- and steroid hormone-producing cells under disease conditions^22,30–32^. In adipose tissue, LAMs possess dual functions. At homeostasis, LAMs facilitate tissue homeostasis by clearing cell debris from dead adipocytes and contribute to the remodeling of the extracellular matrix^33,34^. In contrast, under obesity conditions, the LAM population expands drastically, and they acquire a pro-inflammatory phenotype^35–37^. This alteration exacerbates local inflammation and insulin resistance, thereby promoting systemic dysregulation of metabolism. In the liver, LAMs are essential to the development and progression of NAFLD^15,38,39^. These cells attempt to clear and digest excess lipid, through the upregulation of cholesterol efflux and lipolysis pathways, but in the setting of obesity and atherosclerosis these cells have been associated with excessive lipid overload and dysfunction. This lipid deposition causes the release of inflammatory mediators and promotes hepatic inflammation. Chronic hepatic inflammation eventually leads to liver fibrosis and irreversible damage. LAMs are also essential for the progression of atherosclerosis, a chronic inflammatory disease of the arterial wall. In the arterial intima, macrophages take up oxidized low-density lipoproteins (oxLDL) and become dysfunctional once transformed into lipid-laden foamy macrophages^17,40–44^.

Accumulation of foam cells in the intima ultimately drives atherosclerotic plaque progression. This process poses risks to cardiovascular health.

The molecular mechanisms underlying the formation and function of LAMs are not fully elucidated, but it is proposed that adipose LAMs depend on Trem2 for LAM programming^45^. In recent atherosclerosis studies, conditional deletion of Trem2 in myeloid cells results in slowed atherosclerotic plaque growth^43^. Inversely, therapeutically agonizing Trem2 with an antibody stabilized plaque by promoting collagen accumulation and enhancing LAM survival.

Furthermore, the expression of lipid-handling enzymes in LAMs, including LPL, facilitates their capability to perform lipolysis to breakdown and utilize the internalized lipids. Targeting LAMs offers potential therapeutic benefits, and in fact, many of these approaches are underway in the cancer field. For example, peroxisome proliferator-activated receptor (PPAR) inhibition is used to regulate lipid metabolism in lipid and tumor associated macrophages^46^. Fatty acid binding protein (FABP) inhibition intervenes in lipid transport proteins to reduce lipid handling in LAMs^47^. Similar inhibition approaches have also been developed to target other molecules regulating lipid uptake and accumulation in macrophages, including CD36, a fatty acid receptor and transporter on macrophages^48^,and ABC transporters, which play key roles in the efflux of cholesterol^14,49,50^. Overall, the field of LAM biology still lacks a universal documentation of LAM feature profile to potentially facilitate further therapeutic discovery.

In this study, we aimed to profile representative LAM signatures across chronic inflammatory diseases in mice, using scRNAseq data and integration approaches. We integrated macrophages from multiple tissues and diseases to identify gene programs that are conserved by LAMs across organs, as well as define genes that are unique to LAMs in specific disease settings. Integration analysis of published human scRNAseq data revealed that LAMs are transcriptionally conserved between human and mouse. Overall, these data can be used as a valuable resource for defining LAM subsets, as well as providing insight into tissue-restricted function of LAMs.

## Methods

### Animal maintenance

Ldlr^-/-^ mice were purchased from the Jackson Laboratory (Jax 002207). Animals were housed in the University of Minnesota (UMN) Research Animal Resources (RAR) facility under 12-hour dark and light cycle and specific pathogen-free conditions. Mice had unrestricted access to food and water. High fat diet (HFD) used in this study was purchased from Envigo Teklad (TD.88137, adjusted calories diet, 42% fat). Mice started on HFD at 7 weeks of age and stayed on HFD for 16 weeks. All experiments performed in this study were approved and performed in accordance with the UMN Institutional animal care and use committee (IACUC).

### scRNAseq analysis

#### Preprocessingh

Single cell RNA sequencing (scRNA-seq) datasets were downloaded from gene expression omnibus (GEO) database (Table 1). Data was first loaded and transformed to Seurat object using *Read10X* and *CreateSeuratObject* functions from the Seurat v4.2.1 package. We skipped preprocessing procedures on datasets that were annotated and processed by the original authors. Otherwise, datasets were filtered on cell mitochondrial content, where cells with over 25% mitochondrial transcription were removed. Normalization and scaling were performed using the *NormalizeData* and *ScaleData* functions. Doublets were detected and removed using the DoubletFinder v2.0.3 package.

**Table 1.** Mouse scRNAseq datasets.

| Tissue | Disease | GEO | Authors |
| --- | --- | --- | --- |
| Adrenal gland | Acute/chronic stress | GSE268522 | Xu et al.<sup>87</sup> |
| Adipose | Obesity | GSE182233 | Cottam et al.<sup>88</sup> |
| Liver | NFDLD | GSE166504 | Su et al.<sup>89</sup> |
| Aorta | Atherosclerosis | GSE256189 | Patterson, Xu, Hillman et al.<sup>43</sup> |
| Aorta | Atherosclerosis | GSE116240 | Kim et al.<sup>90</sup> |
| Heart | MI | GSE163465 | Jin et al.<sup>91</sup> |
| Heart | HFpEF | GSE236585 | Li et al.<sup>92</sup> |

#### Data integration

All datasets were first merged as one meta dataset using the *merge* function.

*FindVariableFeatures*, and *RunPCA* functions were subsequently performed. A total of 33947 cells were generated in the mouse metadata, and human data consisted of 11767 cells. To correct batch effects among datasets, *RunHarmony* function from the Harmony v0.1.0 package was applied to the merged object. Next, cells were projected to uniform manifold approximation and projection (UMAP) embedding using the *RunUMAP* function from Seurat, with the “reduction” command set to “harmony”. Determination of significant dimensions was facilitated by an elbow plot generated using the Seurat *ElbowPlot*. *FindNeighbors* and *FindClusters* functions were used to calculate clusters. For LAM reclustering, the LAM population was first identified according to the transcriptional profile of the cluster. Then, the cluster corresponding to the LAMs was extracted. A total of 5941 LAMs were selected. The data integration process described above was repeated on the isolated LAM cluster.

#### LAM feature identification

*FindAllMarkers* function was first used to generate differentially expressed genes for each cluster. The cluster featuring lipid associated genes was annotated as LAM cluster. Next, *FindAllMarkers* was performed on each dataset between the potential LAM cluster and all other clusters to generate LAM-representing features for each tissue and condition. Next, we found the overlapping features across all tissues. In mouse, 14 features were identified. In human, 15 features were identified. Merging of all LAM features was achieved using the *addModuleScore* function and named “LAM score”.

#### Trajectory

Pseudotime trajectory analysis was supported by the Monocle v2.18.0 and Monocle3 v0.2.3.3 packages. First, *as.cell_data_set* function from the SeuratWrapper v0.3.0 package was used to transform Seurat object to Monocle 3 cell_data_set. Root of the pseudotime trajectory was set to classical monocytes identified by transcriptional features, including *Ly6c2*. Next, *cluster_cells, lean_graph,* and *order_cells* were subsequently performed.

#### KEGG pathway analysis

*FindMarkers* function was used to generate the gene background for LAM cluster, with logfc.threshold and min.pct argument set to 0. Functions from the fgsea v 1.18.0 package were used to performed pathway analysis. KEGG pathway database was imported from the msigdbr v7.2.1 package using the *msigdbr* function.

##### Flow cytometry

Freshly harvested adrenal gland, liver, aorta, and white adipose were finely minced. Adrenal glands were enzymatically dissociated using Liberase DH (Roche 5401054001, 2.5mg/mL in double:200-distilled water) at 10% dilution for 35 minutes. Liver, aorta, and white adipose were processed in 1.5mg/mL collagenase A (COLLARO 10103578001) for 35 minutes. Dissociation procedure was performed at 37°C in orbital shaker at 800 round per minute. Dissociated single cell suspensions were next filtered through 100μm nylon mesh (McMaster Carr) to remove debris. Cell solutions next underwent centrifuging at 1450 round per minute for 5 minutes. For antibody staining, all cells were stained at 1:200 diluted flow antibodies (Table 3) in FACS buffer at 4°C for 25 minutes. Cells were washed twice in FACS buffer post staining, and data was collected using the BD LSRFortessa instrument maintained at the flow cytometry core facility at the University of Minnesota.

##### Statistics

Flow cytometry graphs were made in Flowjo software. Graphpad Prism was used to generate bar graphs and perform statistical analysis, where comparisons of group more than 2 were performed by ANOVA. Statistical significance were indicated using: ns (P>0.05), * (P<0.05), ** (P<0.01),*** (P<0.001) and **** (P<0.0001). Differential expression was based on the non-parametric Wilcoxon rank sum text, supported by the Seurat package. P-value adjustment shown in this study used the Bonferroni correction method.

## Data availability

Previously published scRNAseq datasets involved in this study are publicly available. Further analysis of data and code scripts will be made available upon request.

## Results

### scRNAseq integration reveals macrophage heterogeneity in response to chronic inflammation

Macrophages have been appreciated for their functions in disease pathogenesis and resolution. Shared macrophage subpopulations are recognized to emerge following injury or chronic inflammation^1,51,52^, but the transcriptional profiles of these cells have not been thoroughly investigated. To computationally investigate the commonalities and differences of macrophage subsets from different tissues in response to metabolic dysregulation or chronic inflammation, we integrated scRNAseq data from atherosclerotic aorta, livers from nonalcoholic fatty liver disease (NAFLD), hearts with heart failure with preserved ejection fraction (HFpEF), white adipose tissue (WAT) under high-fat diet fed obesity conditions, and adrenal glands under stress conditions (Table 1, Fig 1A). We also included hearts bearing myocardial infarction to investigate macrophages responses to more acute sterile inflammation. Since we were particularly interested in cells under pathophysiological conditions, we did not include cells from healthy mice in the integration analysis. We first performed quality control to remove low quality cells based on low read depth and high mitochondria content. Next, monocyte and macrophages (MoMACs) were extracted from the datasets according to the original cell type annotation by the authors, after which post-processed cells underwent integration using *Harmony* and further downstream analysis (Fig 1A). Clusters of monocytes and macrophages were subsequently projected to uniform manifold approximation (UMAP) space, where 9 clusters were identified (Fig 1B). All 9 clusters were detected in different tissues and conditions, with cluster 0 and 1 being the dominant populations (Fig 1C, D, E). The high uniformity of clusters between tissues may suggest MoMACs potentially undergo similar transcriptional programing in response to metabolic dysregulation or tissue injury, regardless of the homing site (Fig 1E).

**Fig 1.**
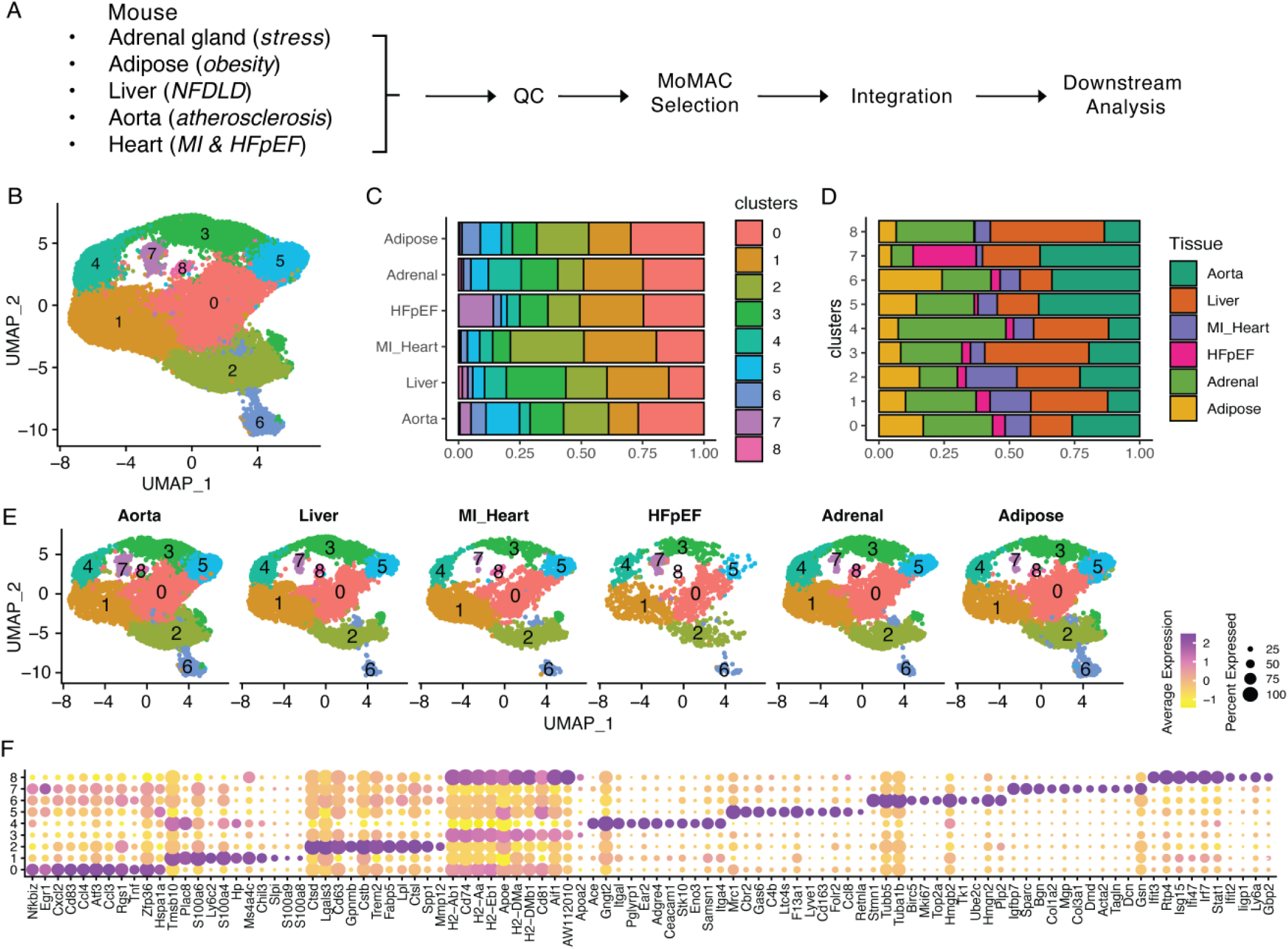
Integration of mouse monocytes and macrophages across tissue. **A.** Workflow for the design of the integration study. **B**. Clusters of monocytes and macrophages shown in UMAP embedding. **C.** Cluster composition of each tissue and condition. **D**. Tissue and condition composition for each cluster. **E**. UMAP cluster split by tissue and conditions. **F**. Differentially expressed genes that represent each cluster, shown in a dot plot.

Next, differential expression analysis was performed to show the leading features of each cell cluster. Cluster 0 appeared to be classically activated macrophages as it yielded high expression of *Tnf* and pro-inflammatory chemokines, such as *Ccl4*, *Ccl3* and *Cxcl2* (Fig 1F).

Cluster 1 appeared high for *Ly6c2* and *Hp*, and thus identified as classical monocytes. Notably, cluster 2 exhibited prominent expression of lipid sensing molecules, such as Trem2 and Lpl, indicating these were LAMs. Major histocompatibility complex II (MHC-II) molecules, *H2-Ab1* and *H2-Eb1*, and *CD74* are high for cluster 3, suggesting these cells are prone to antigen presentation. Cluster 4 was distinguished by *Ace* and *Adgre4*. Interestingly, we were able to identify cluster 5 as Lyve1-resident macrophages, that also co-expressed *Cd163*, *Mrc1* and *Folr2,* which has been documented in previous study^8^ . Cluster 6 showed high expression of *Top2a* and *Mki67*, therefore distinguished as proliferating macrophages. According to our previous study, there was a smooth muscle cell derived macrophage population in atherosclerotic lesions that showed prominent expression of *Acta2*^43^. Rather surprisingly, this subset of cells indeed appeared across all tissues in this integration analysis in cluster 7. In addition, collagen associated genes, *Col1a2* and *Col1a3*, were differentially expressed in this cluster, suggesting these macrophages may be homeostatic by promoting collagen deposition. Interesting, cluster 8 was distinguished by *Ifit2* and *Ifit3* expression, that are linked to type I interferon signaling, suggesting this population potentially responds to inflammatory stimuli^53,54^. Overall, this integration analysis across disease and tissues suggested that macrophages may possess a conserved transcriptional response to cope with tissue injury.

### LAMs transcriptional signatures are conserved across tissue and condition

Studies have shown that LAMs are vital participants in metabolic homeostasis and regulation^11,55–57^. However, it is well recognized that LAMs play various roles in different tissues^39,58–62^. Whether LAMs from different tissue origins are transcriptionally conserved in mice is not well understood. To address this question, we first identified LAMs from the integrated meta-dataset. Since *Trem2* and *Lpl* are well characterized LAM associated genes^13,14^, we used these two molecules to first identify cluster 2 as the LAM population (Fig 1F). Next, differential expression analysis between cluster 2 and all other cells was performed on each tissue independently. This step generated a set of genes for each tissue that represent LAM features in that tissue. This gene pool was integrated together, and we identified genes that were uniquely expressed by LAMs and their enrichment was shared by all tissues. 14 overlapping genes were identified, including *Cd63, Cd68, Cstb, Ctsb, Ctsd, Fabp5, Gpnmb, Lgals3, Lipa, Lpl, Pld3, Plin2, Spp1* and *Trem2* (Fig 2A). Merging these genes into a pathway feature plot, resulted in co-expression in cluster 2 (Fig 2B, 2C), validating that cluster 2 is a conserved LAM cluster.

**Fig 2.**
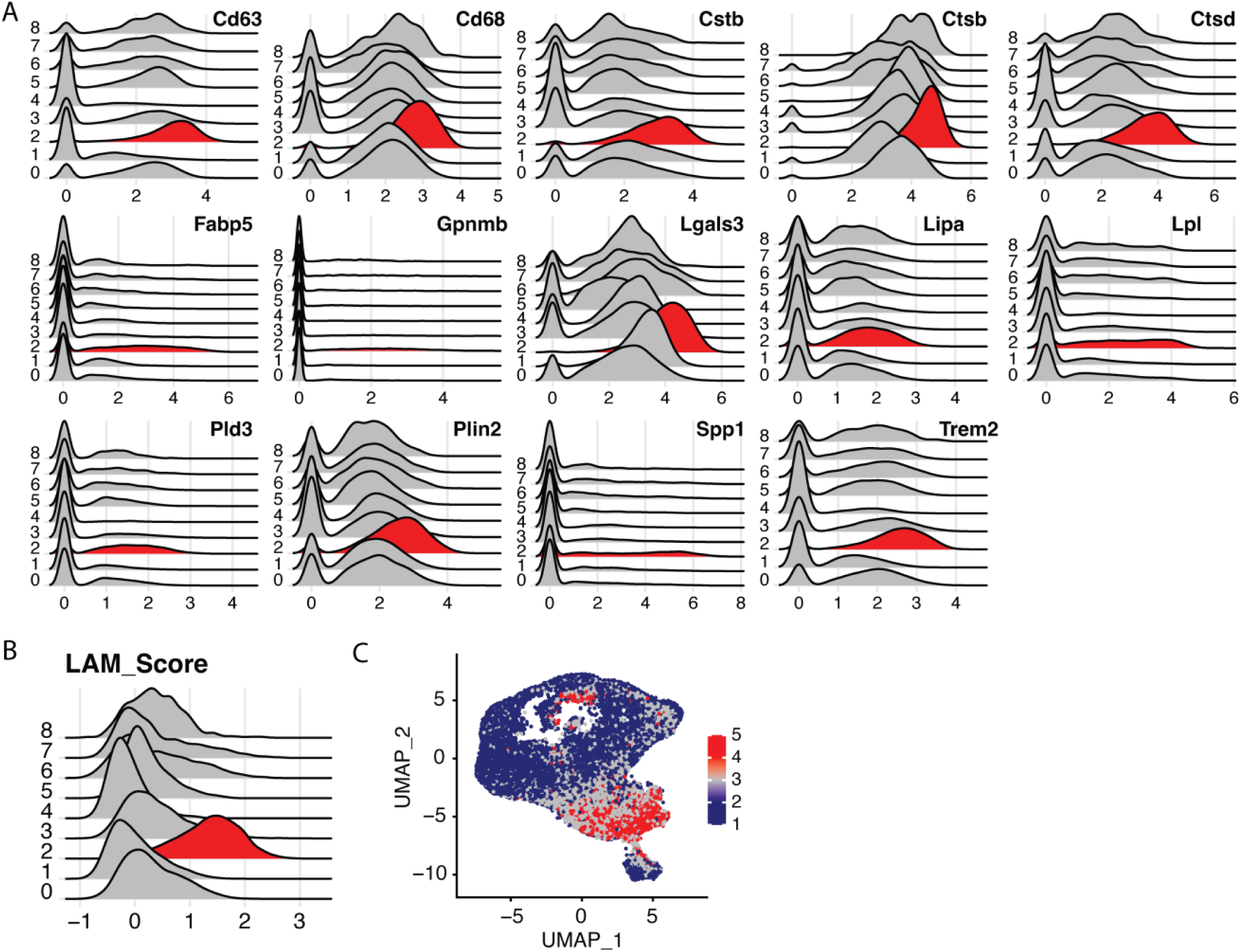
Conserved lipid-associated macrophage features in mouse. **A.** Conserved 14 features that represent LAMs in different tissue and condition. **B**. Features merged and named LAM score. **C**. LAM score expression in UMAP space.

Of these 14 features, *Cd63, Cd68, Ctsb* and *Ctsd* are associated with macrophage lysosomal functions^65,66,66^. Lipid sensing and signaling also appeared to be highlighted in LAMs as *Fabp5, Lipa, Lpl, Trem2* and *Pld3* are highly expressed^27,63,64^. In addition, the expression of *Gpnmb* and *Spp1* may indicate LAMs participate in the regulation of inflammation^67–69^. Next, KEGG pathway analysis was performed to reflect biological and cellular functions of genes.

Notably, lipid utilization was reflected by PPAR signaling, steroid, and glycerolipid metabolism (Sup 1A). LAMs widely present lipid metabolism associated homeostatic characteristics as oxidative phosphorylation and lysosomal associated activities, including both proteosome and lysosome activation, are upregulated (Sup 1A). Together, the data support that LAMs possess complex functions to regulate lipid homeostasis, a feature that is conserved across tissues.

Furthermore, we investigated the potential developmental trajectory of LAMs from different tissue origins and conditions. According to previous studies in atherosclerosis, macrophages from tissue-resident and monocytic origins share a similar transcriptional profile^70,71^. Under high-fat diet induced obesity, WAT LAMs primarily derive from monocytic lineage^72,73^, but whether LAMs from other tissue niches follow the same differentiation pattern remains unclear. We hypothesized that the majority of LAMs arise from circulating monocytes to sustain local populations in different tissues during disease. We took a computational approach by inferring pseudotime trajectory to each dataset to test this hypothesis. Cluster 1, classical monocytes, was set as the root of pseudotime trajectory. We observed a clear trend where pseudotime ends in cluster 2, LAM, and cluster 3, MHC-II high. This pattern was observed in all tissues and conditions, suggesting LAMs across tissue may depend, in part, on monocyte precursors (Sup 1B).

#### LAMs exhibit tissue specific characteristics

Despite the shared features across tissues, we next sought to investigate tissue-specific markers that may represent LAM population heterogeneity across tissues. First, we performed re-clustering of MoMAC cluster 2. 4 LAM subclusters were generated (Fig 3A). Subcluster 0 appeared to be the dominant cluster. Interestingly LAM cluster 3 seemed to be unique to mouse aorta bearing atherosclerosis, and it was completely missing from other conditions and tissues (Fig 3B). This was shown more clearly in cluster split by tissue, where cluster 3 only appeared in the aorta dataset (Fig 3C, 3D). Next, differential expression was performed to calculate features representing each subcluster. The canonical LAM markers, such as *Lpl, Mmp12,* and *Spp1,* were expressed by all cells, although the average expression for these features vary (Fig 3E).

**Fig 3.**
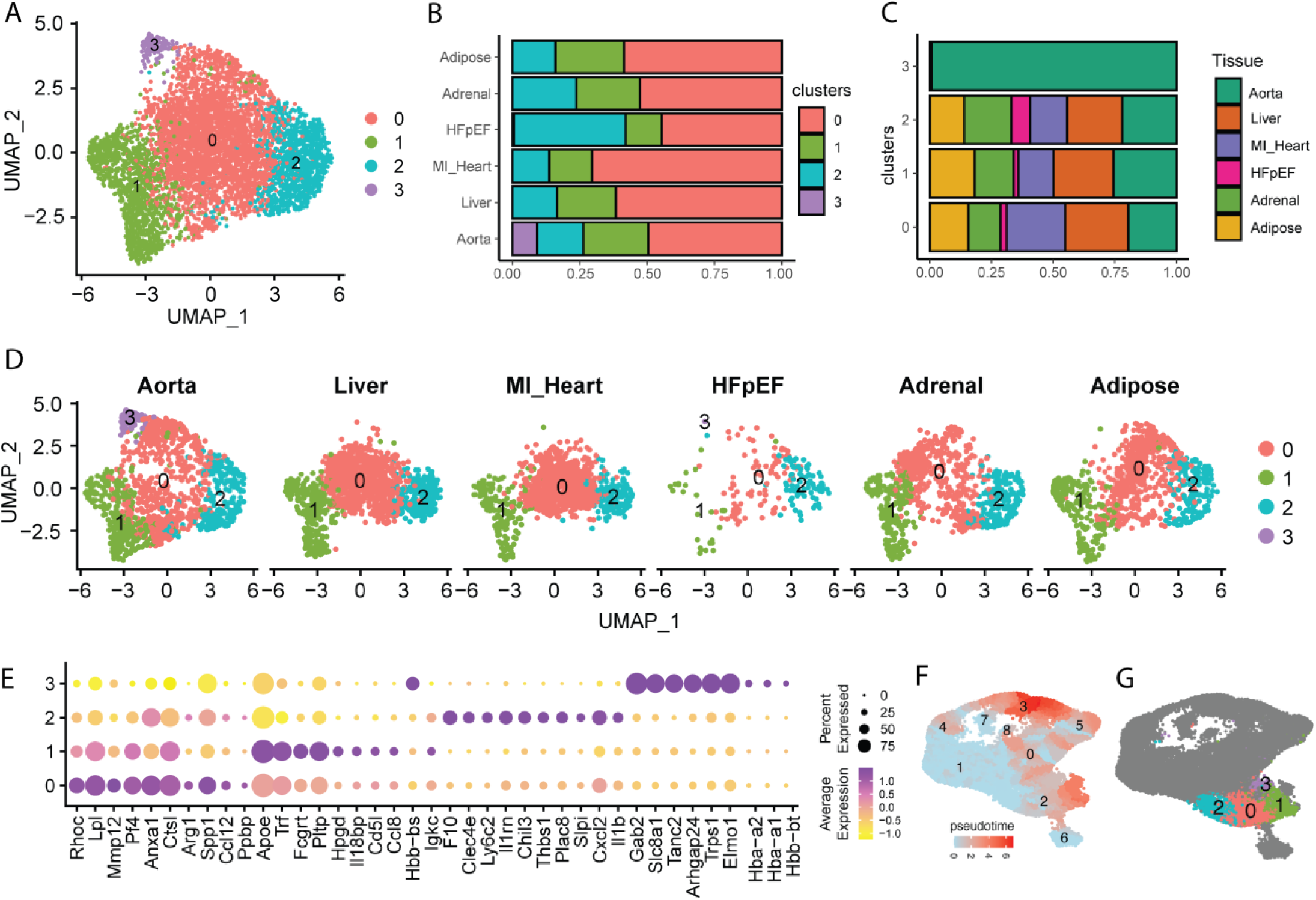
Reclustering analysis revealed high heterogeneity of LAM. **A.** UMAP embedding showing 4 LAM subclusters. **B**. Cluster composition for each tissue. **C**. Tissue/condition composition for each LAM cluster. **D**. UMAP cluster split by tissue. **E**. Top 10 signature genes for each LAM cluster. **F**. Pseudotime trajectory involving all monocytes and macrophage in UMAP space. **G**. LAM subclusters shown in previous total MoMAC UMAP space.

Subcluster 0 and 1 were particularly high for these markers. Subcluster 2 showed relatively high expression of classical monocyte markers, such as *Ly6c2*, and therefore identified as newly differentiated LAMs deriving from recently recruited monocytes. Notably, subcluster 3 was distinguished by high *Gab2, Slc8a1* and *Tanc2*, which was specific to aorta. To computationally develop a more defined differentiation trajectory with the LAM subpopulations, we reintroduced pseudotime to the new clusters. Interestingly, subcluster 2 was upstream of trajectory pseudotime, suggesting this is the population that first to differentiate and adopt LAM phenotype (Fig 3F, 3G). Since subcluster 2 expresses lipid associated genes while maintaining genes such as *Ly6c2*, reflecting a monocyte phenotype, we argued that this stage of cells could be indicative of early events leading to LAM differentiation. To investigate this idea, we performed differential gene expression analysis between monocyte cluster 1 and LAM subcluster 2 (Sup 2A). Notably, we observed prominent upregulation of LAM features in subcluster 2 including *Fabp5, Gpnmb, Ctsd, Cd63, Cstb, Lgals3, Plin2, Cd68, Trem2, Cd9, Spp1, Lpl,* and *Mmp12.* Of these markers, *Fabp5, Gpnmb, Spp1,* and *Mmp12* were at a more significantly upregulated state (Sup 2B), suggesting LAMs may initially turn on these genes during differentiation toward mature LAMs. Subcluster 0 subsequently emerged from cluster 2. Compared with cluster 1, cluster 0 showed lower expression of *Apoe*, which may suggest that this population has already undergone LAM transcriptional programming but is not at a fully mature differentiation state (Fig 3F, 3G). This reclustering approach revealed unique heterogeneity and composition of LAMs during disease and proposes the co-existence of several LAM populations in tissues, likely with common origins from monocyte precursors.

Next, we specifically investigated the differences of LAMs across tissues. Although, the majority of differentially expressed genes (DEGs) are not tissue-specific, aortic and HFpEF LAMs exhibited the most segregated gene expression patterns, where 447 and 500 features were differentially expressed, respectively (Fig 4A). Liver, MI heart, WAT, and adrenal LAMs showed 97, 71, 240 and 191 DEGs, respectively, suggesting LAMs indeed possess tissue-specific characteristics (Fig 4A). Aortic LAMs, in particular, showed high expression of *Abca1*, consistent with aortic LAMs promoting cholesterol efflux pathways during atherosclerosis progression (Fig 4B, 4C). *Lyz1* was exclusively expressed in liver LAMs. In addition, these cells showed distinguishable *Id3, Ear2,* and *Apoc1* expression (Fig 4B, 4C). As for hearts bearing MI condition, *Arg1* was highly enriched. Interestingly, chemokines associated with acute inflammation and monocytes recruitment, including *Ccl2, Ccl12 and Ccl7,* were also strongly upregulated. This may suggest LAMs in MI heart potentially participate in inflammatory response by contributing to monocyte recruitment. Interestingly, HFpEF LAMs were defined by *Ptgs2* expression. It has been shown that macrophage expression of *Ptgs2* mediates proinflammatory response by driving immune cell infiltration to inflammatory sites^74,75^. It is also involved in the resolution phase of inflammation and contributes to tissue homeostasis. This observation supports the idea that LAMs in HFpEF are indeed a regulator of chronic disease.

**Fig 4.**
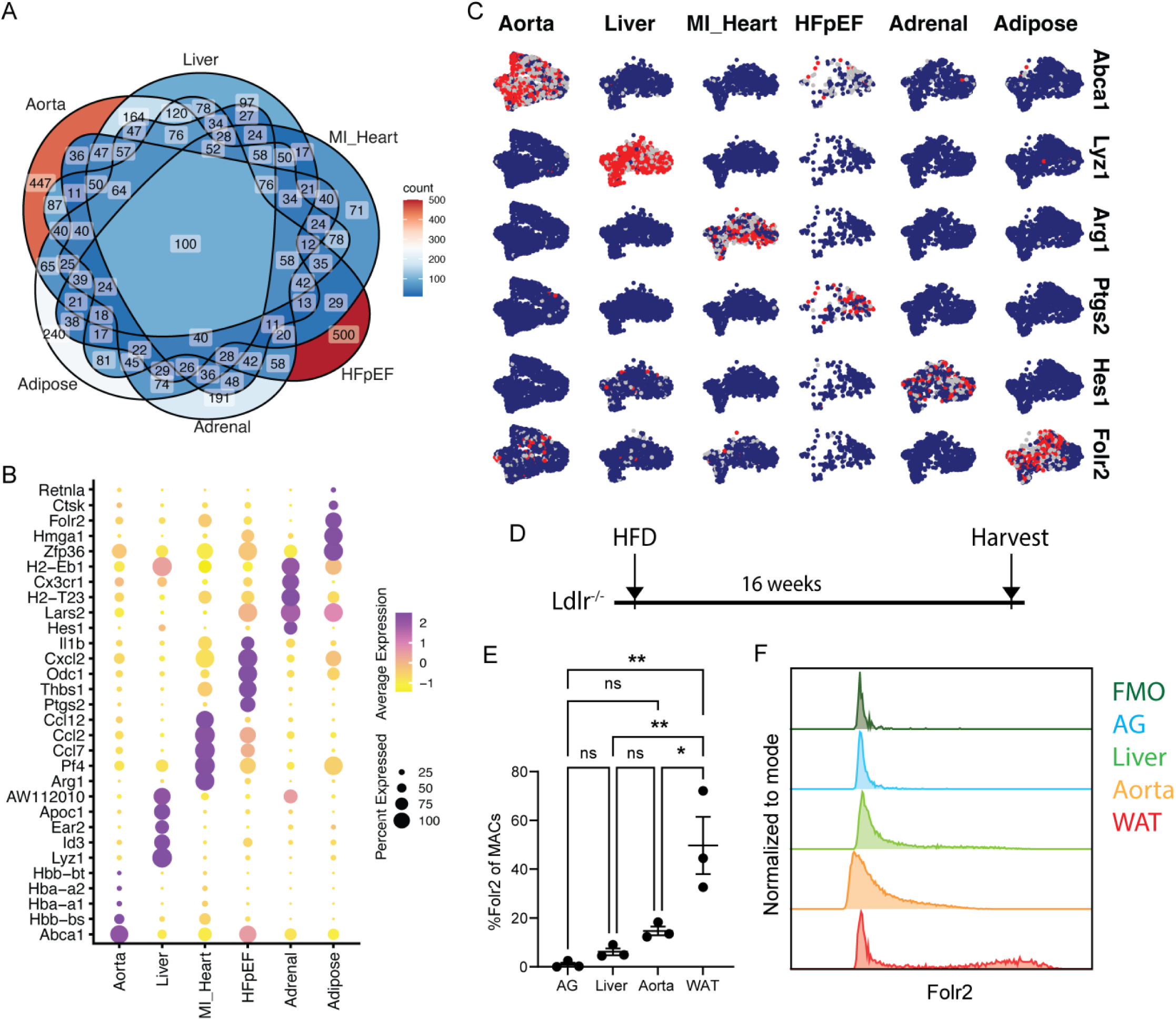
LAMs exhibit transcriptional tissue specificity. **A.** Venn diagram shown overlapping and unique gene count between tissues in LAM. **B**. Top 5 differentially expressed genes that represent each tissue. **C**. Top representative feature for each tissue. **D**. Schematic of experimental design showing induction of metabolic dysregulation in mouse model. **E**. Quantification of Folr2+ macrophages in tissue (n=3 for each group). **F**. Histogram showing Folr2 expression in different tissues.

Furthermore, adrenal LAMs exclusively express *Hes1*, which is considered an anti-inflammatory gene^76^, suggesting adrenal LAMs may utilize *Hes1* to dampen inflammation by limiting immune cell infiltration (Fig 4B, 4C). Lastly, adipose LAMs showed high expression of *Folr2* (Fig 4B, 4C), which in most tissues is restricted to a subpopulation of tissue resident macrophages^8^. In summary, differential analysis of LAMs across tissues has identified tissue-specific expression patterns that are likely associated with specialized functions in these pathologic responses.

To experimentally validate our observations regarding tissue specificity of LAMs, we utilized Ldlr^-/-^ mice fed on high fat diet (HFD) for 16 weeks (Fig 4D). This diet induces systemic elevation of cholesterol, therefore promoting atherosclerosis, metabolic dysregulation, and disease progression. Flow cytometry analysis was used to quantify target molecules. Since only a limited number of LAM genes have commercially available antibodies for flow cytometry, we were restricted to the targets we could interrogate. Here, we aimed to measure Folr2 enrichment in the aorta, as well as the adrenal gland, liver, and WAT, as antibodies targeting Folr2 are widely available. We defined macrophages in tissues as sharing expression of F4/80 and CD11b. Total macrophages were then examined for significant Folr2 expression, where positivity in WAT macrophages, where about 40% of WAT macrophages were classified positive for Folr2 (Fig 4E, 4F, Sup 3). Consistent with transcriptional data, aortic macrophage Folr2 expression ranked second after WAT, while adrenal gland and liver macrophages showed very little Folr2 expression (Fig 4E, 4F). In conclusion, our computational and experimental approaches suggest LAM indeed possess tissue specificity despite their overlapping core LAM transcriptional profile.

#### LAMs signatures are conserved in human

Despite the number of studies reporting LAMs in mouse, only a few studies have directly investigated gene programing for LAM between species^77,72,70^. To investigate this idea, we integrated monocytes and macrophages from four human studies, including samples from obese WAT, liver cirrhosis, atherosclerotic plaque, and dilated cardiomyopathy (DCM) biopsy (Fig 5A, Table 2). Monocyte and macrophages were first isolated according to the author’s annotations. Next, we used CD14 expression to validate the annotation and purify the MoMAC population, followed by integration of individual datasets and downstream analysis. Clustering of the integrated dataset resulted in 7 MoMAC clusters (Fig 5B). Although the overall composition of clusters was similar between tissues, adipose tissue showed a lower proportion of cluster 2 and enrichment of cluster 1 and 5 (Fig 5C). Interestingly, the heart exhibited a unique cluster 6 which was absent in the other tissues (Fig 5C, 5D, 5E). Differential analysis was performed to generate transcriptional features for each cluster. Cluster 0 and 6 showed a high expression of LYVE1, suggesting this was a tissue resident macrophage population (Fig 5F). Overlapping features between 0 and 6 also included C1QA, PLTP, DAB2, TXNIP, and F13A1. SEPP1 was unique to cluster 0, and ATP6V0C and NME2 were unique to cluster 6. Cluster 1 was distinguished by SPP1, FABP4, APOC1, TREM2, and MMP9, likely the LAM population. Cluster 2 exhibited mostly ribosomal genes, suggesting this might be stressed macrophages. Cluster 3 showed monocyte features like S100A8, S100A9 and S100A12. Interestingly, we observed immunoglobulin associated gene expression, such as IGHM, IGLC2 and IGLC3 in cluster 4.

**Fig 5.**
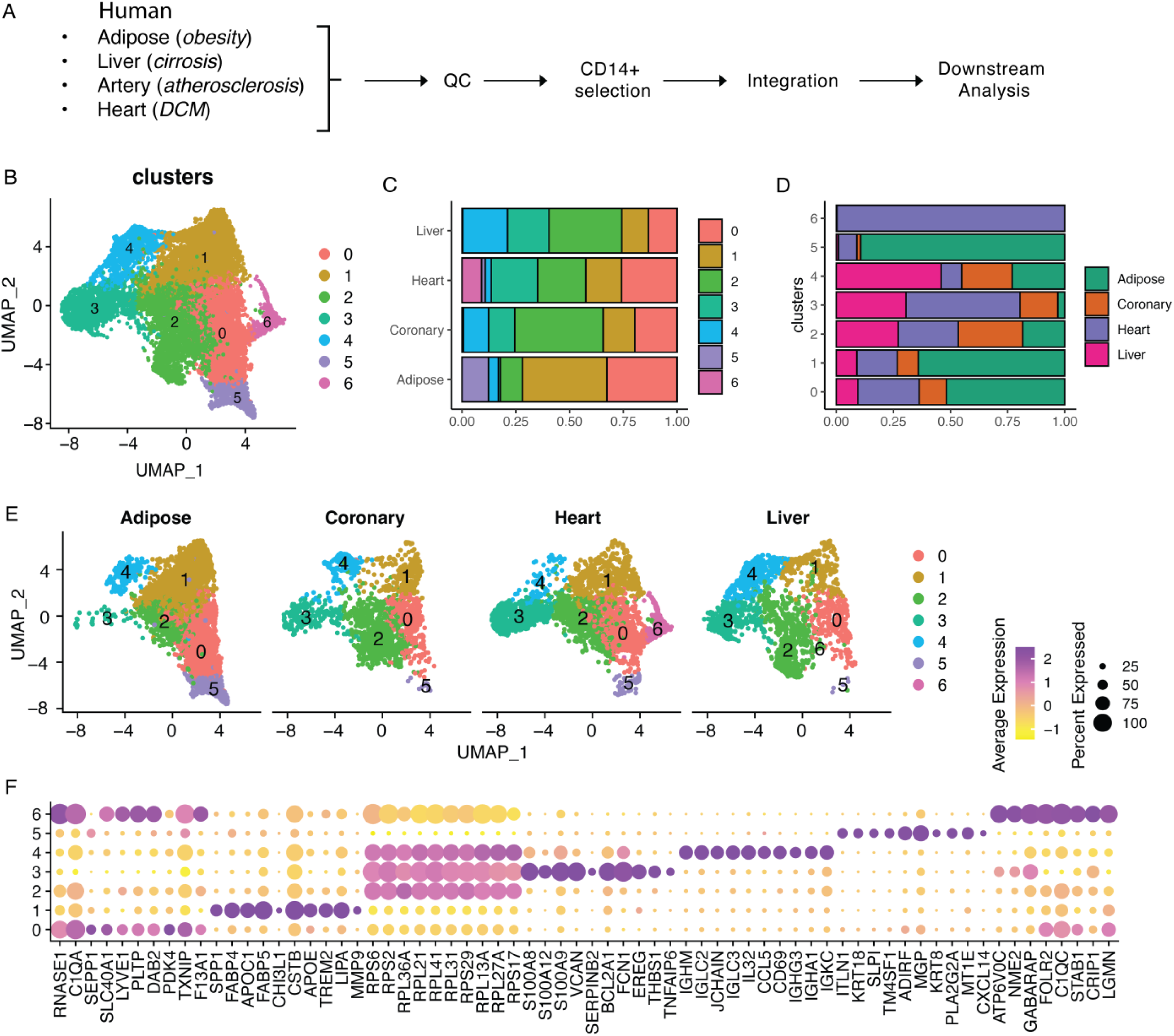
Integration of human scRNAseq dataset across tissue. **A.** Schematic workflow for the integration analysis. **B**. Human MoMACs shown in UMAP space. **C**. Cluster composition for each tissue/condition. **D**. Tissue composition for each cluster. E. UMAP clusters split by tissue. **F**. Top 10 differentially expressed genes shown in dot plot.

**Table 2.** Human scRNAseq datasets.

| Tissue | Disease | GEO | Authors |
| --- | --- | --- | --- |
| Adipose | Obesity |  | Massier et al.<sup>93</sup> |
| Heart | DCM | GSE183852 | Koenig et al.<sup>94</sup> |
| Liver | Cirrhosis | GSE136103 | Ramachandran et al.<sup>95</sup> |
| Coronary artery | Atherosclerosis | GSE131778 | Wirka et al.<sup>96</sup> |

**Table 3.**
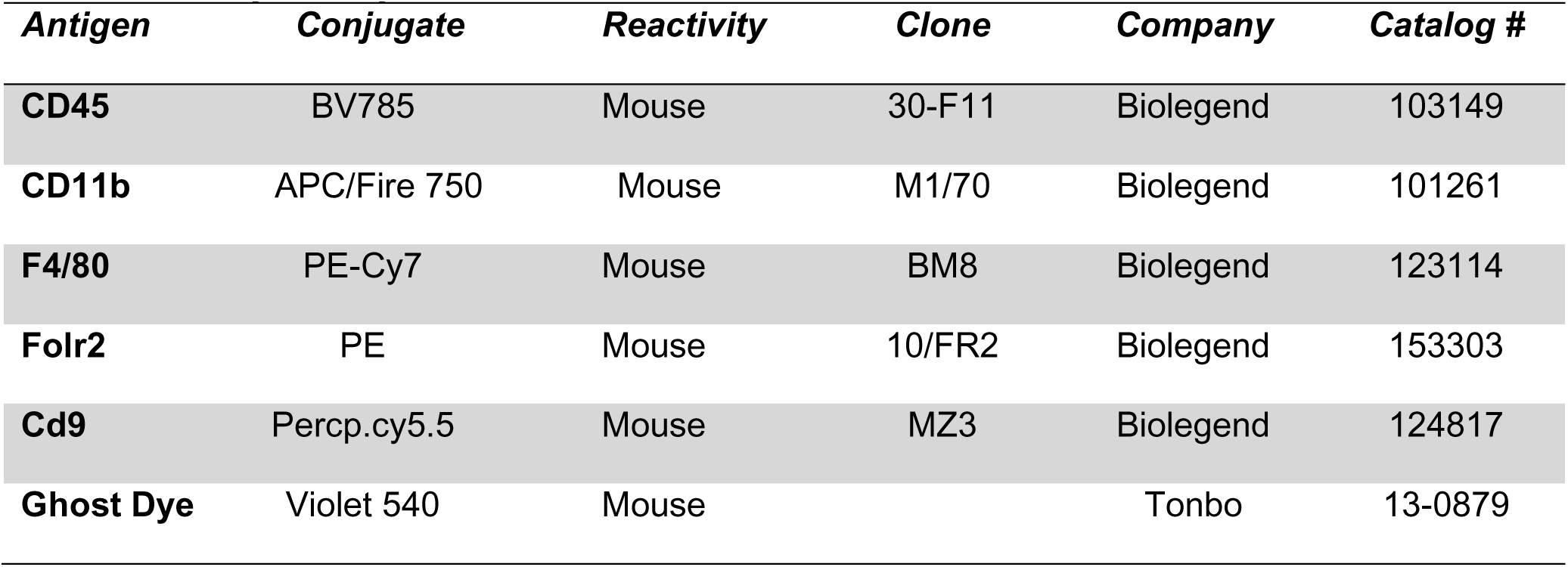
Flow cytometry antibodies.

CCL5 and CD69 were also highly expressed in this cluster, which exhibited a B cell signature. We argued that this might be macrophages being contaminated by B cells. Cluster 5 showed unique expression of ITLN1, KRT18, SLPI, and TM4SF1. Overall, human MoMACs exhibited high heterogeneity as cell clusters differentially expressed unique features. However, consistent with recent studies comparing tissue-resident macrophages between mouse and human, several key overlapping signatures were conserved.

Next, we took a similar approach to our mouse study to generate LAM features in human cells. We first calculated DEGs of cluster 1 for each tissue dataset, as cluster 1 co-expresses multiple lipid associated features, such as TREM2 and LIPA. We aimed to find overlapping genes across all tissues and named it LAM score. A total of 15 genes were generated, including FABP4, FABP5, APOC1, CSTB, SPP1, LIPA, TREM2, CTSD, CTSB, CD9, APOE, LGALS3, ACP5, GPNMB, and PLD3 (Fig 6A, 6B, 6C). Interestingly, most of these genes overlapped with mouse LAM signatures. To address whether LAM programming was conserved between human and mice, we overlaid the 14 mouse LAM consensus genes on the human LAM DEG background. Mouse LAM features were strikingly consistent with that of human (Fig 6D). This phenomenon suggested that core LAM programming is transcriptionally conserved between human and mouse.

**Fig. 6.**
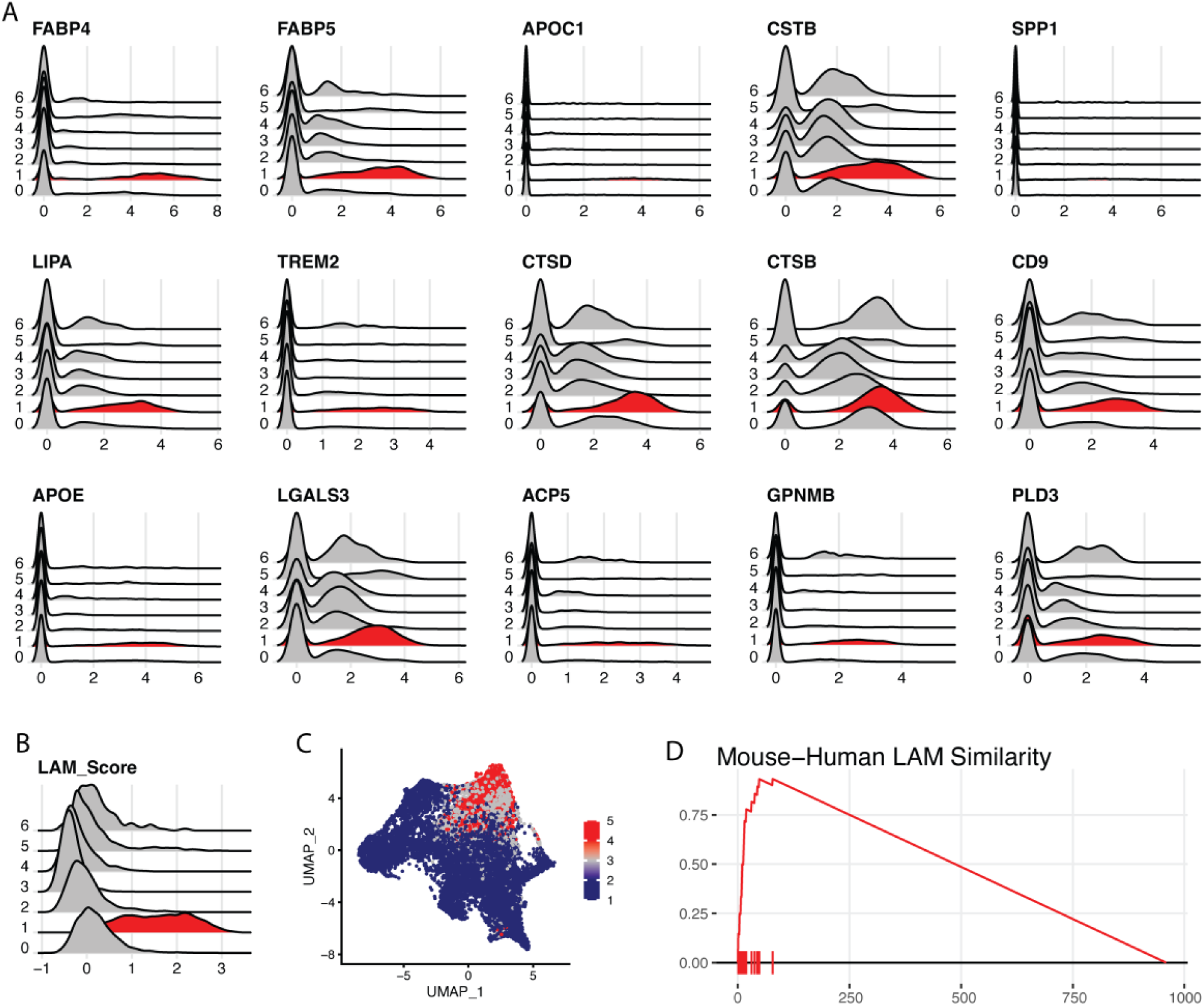
Human and mouse LAMs are transcriptionally conserved. **A.** 15 representative features for human LAMs. **B**. 15 features merged as “LAM score”. **C**. LAM feature expression shown in UMAP space. **D**. Enrichment of mouse LAM feature on human LAM expression background.

Despite the conserved features, a few genes were not shared across human and mouse LAMs. Human LAMs specifically featured APOC1, APOE, ACP5, FABP4, and CD9 (Fig 6A), while mouse exhibit high expressions of *Cd63, Cd68, Lpl, and Plin2* (Fig 2A). Rather surprisingly, mouse LAMs features did not include Cd9, as this gene has been well-documented in previous studies involved in the context of atherosclerosis, liver, and WAT LAMs^18,45,78^.

Absence of Cd9 led us to speculate that some of the tissue may exhibit low expression of the gene. To test this idea, we examined Cd9 expression in each of the tissues. Interestingly, adrenal, aortic, liver, and WAT LAMs showed profound enrichment of Cd9, but it was faintly detected in either MI or HEpEF cardiac LAMs (Sup 4A). Additionally, we performed experimental validation of Cd9 in adrenal gland, aorta, liver, and WAT. As introduced previously (Fig 4D), Ldlr^-/-^ mice and HFD were utilized to disturb lipid metabolism in mice. Consistent with the RNAseq data, Cd9 protein was observed in the 4 tissues after physiological challenge with high fat diet (Sup 4B-3E), suggesting CD9 can indeed be reliable for identifying LAMs in mouse except heart. In conclusion, we observed both conserved and distinct gene signatures of LAMs between conditions and species, underscoring the importance of understanding the tissue-specific LAM signatures in translational study.

## Discussion

This study provides a comprehensive analysis of LAM heterogeneity across tissues in response to sterile inflammation, highlighting both conserved and tissue-specific transcriptional alterations. Using an integration approach of scRNAseq data from multiple metabolic disease models, we documented various macrophage subsets that arose during disease. One subset was found to be LAMs. These findings expand our understanding of how macrophages adapt to metabolic stress and serve as a resource for future studies investigating the function of LAMs in different disease models.

Our analysis revealed that LAMs share common transcriptional features in response to chronic and sterile inflammation. The conservation of LAM transcriptional signatures across different tissues and conditions highlights the role of these cells in maintaining lipid homeostasis and inflammation regulation. The shared set of features across human and mouse suggests that the fundamental mechanisms governing LAM functions are preserved in evolution. Interestingly, all LAM populations show consistent expression of genes such as *Trem2, Lpl, and Fabp5*, across organs, suggesting that LAMs may serve as a critical component in the body’s response to systemic metabolic challenges, regardless of the tissue context. Notably, given the ubiquitous expression of Trem2 by LAMs across tissues and species, therapeutic interventions of Trem2 could represent a promising strategy for modulating dysregulated metabolism in human.

In fact, immunotherapeutic approaches agonizing and antagonizing Trem2 are underway in clinic^79–82^. According to a recent study targeting Trem2 in context of atherosclerosis^43,83^, Trem2 agonist treatment stabilizes atherosclerotic plaque by promoting Trem2-expressing foamy macrophage survival and reducing cell apoptosis. Interestingly, this treatment led to increased collagen deposition in atherosclerotic niche, which also likely contributes to plaque stability. This study not only emphasizes the importance of LAMs in disease progression, but also highlights that therapeutic strategies targeting LAMs are viable to regulate disease outcomes. In the context of breast cancer, Trem2-expressing LAMs have been demonstrated to mediate immune suppression by limiting T cells effector function and proliferative capacity^20,84,85^. Interestingly, anti-Trem2 treatment can indeed enhance anti-PD-1 efficacy and promote immune infiltration in cancer microenvironment, thereby enhancing T cell functions and limiting tumor growth^86^. Together, Trem2 has been shown to possess therapeutic potential. Targeting Trem2+ LAMs could serve as a promising approach against various diseases where LAMs are found.

Despite the conservation of LAM features, our study also highlighted tissue specificity in LAM gene expression, reflecting tissue specific function of LAMs. For instance, aortic LAMs exhibited high expression of *Abca1*, which is associated with cholesterol efflux. Heart-resident LAMs in both myocardial infarction and HFpEF conditions upregulated inflammatory cytokines such as *Ccl2* and *Il1b*, suggesting LAMs may be vital in promoting inflammatory response in the context of the heart. This phenomenon indicates that while LAMs share a core set of functions, their specific roles may be modulated by the local tissue environment, contributing to the unique pathophysiological outcomes observed in different metabolic diseases.

In conclusion, our integrative analysis of mouse and human scRNAseq data offers a valuable reference for the study of lipid-associated macrophages (LAMs), potentially serving as a foundation for future research. We emphasize the significant heterogeneity of LAMs in both mouse and human models, with LAMs displaying conserved transcriptional features across different tissues. Despite these conserved characteristics, LAMs also exhibit tissue-specific transcriptional profiles, which we have validated through experimental approaches. Importantly, the strong overlap in LAM features between mouse and human underscores the utility of mouse models in translational studies, particularly for testing therapeutics aimed at targeting LAMs in human diseases. Overall, our findings highlight the potential for leveraging conserved LAM signatures in both species to develop therapeutic strategies that can address a range of LAM-related conditions across different tissues and diseases.

## Author Contributions

YX, SI, and JWW conceived and designed the project. YX, HH, and MC conducted the experiments. YX conducted the flow cytometry and RNA sequencing data analysis. YX and JWW wrote, and all authors participated in editing the manuscript.

## Acknowledgements & Support

This study was supported by National Institute of Health (NIH) R01 AI165553 (JWW) and NIH R01 HL16683. Thanks to the UMN Flow Cytometry Resource (UFCR) and UMN Minnesota Supercomputing Institute (MSI) for support and assistance in data acquisition and analysis.

---

**Sup Fig 1.**
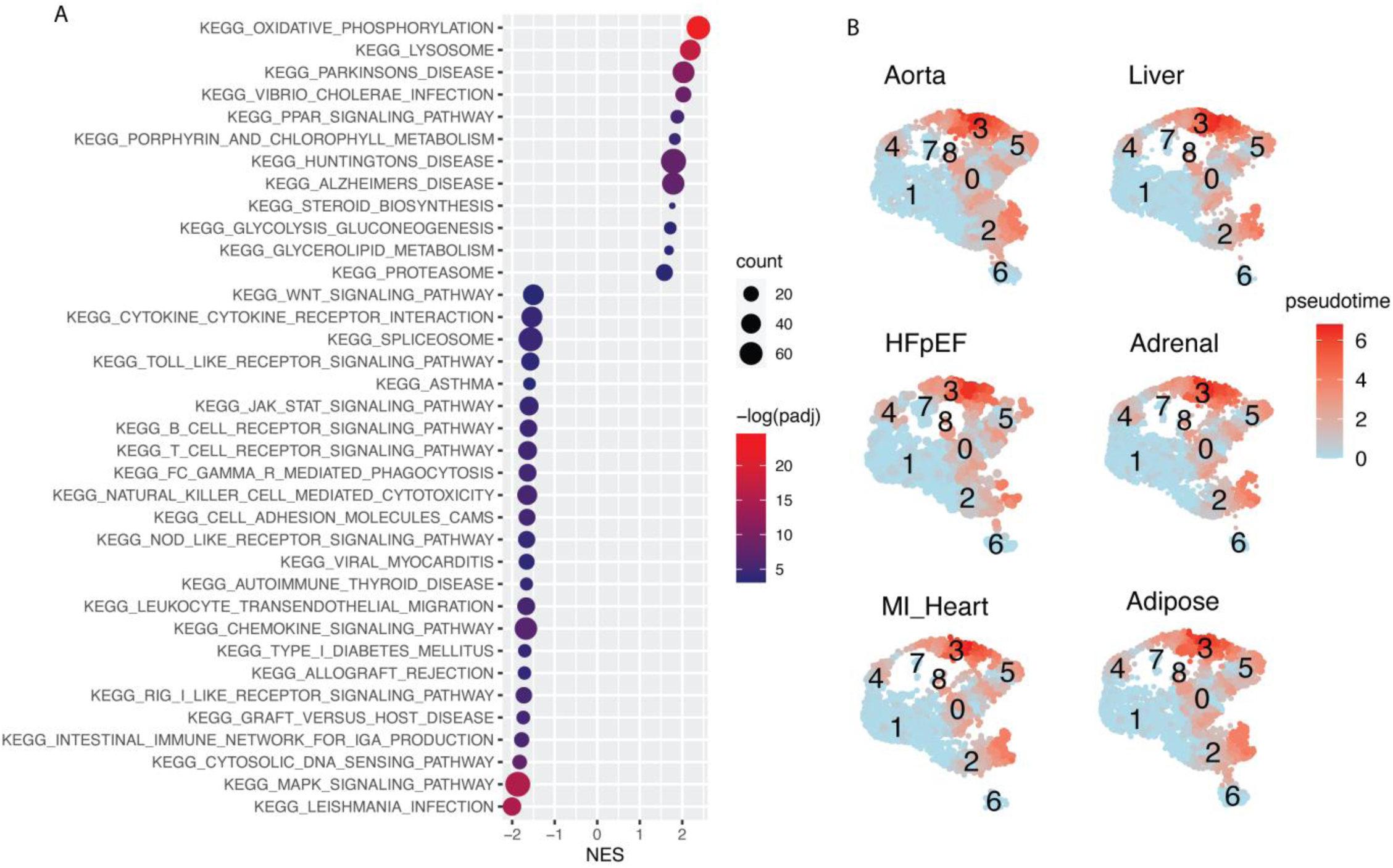
Pathway and pseudotime trajectory analysis of LAM in mouse. **A.** KEGG pathway analysis showing enrichment of functional pathway associated with LAMs. NES: normalized enrichment score. Count: number of gene in the corresponding pathway. -log(padj): log10 transformed adjusted p value. **B**. Pseudotime trajectory of mouse LAMs.

**Sup Fig 2.**
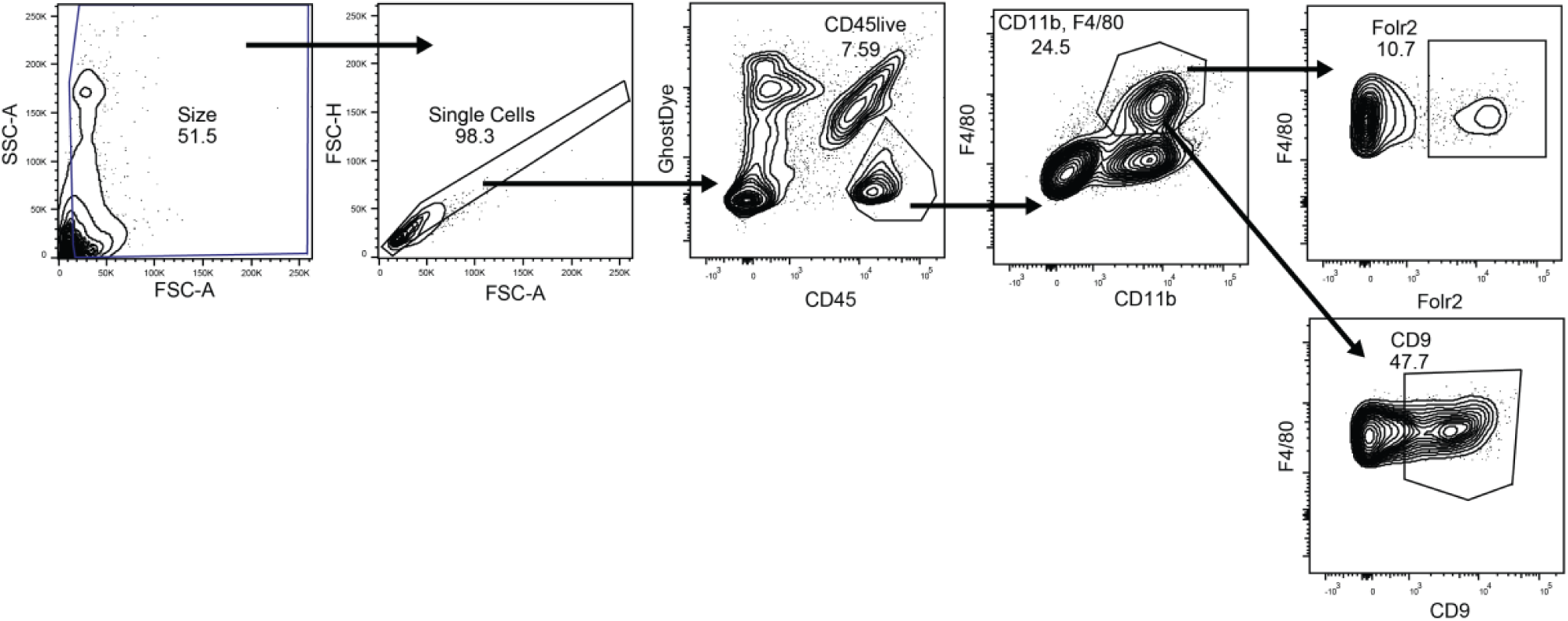
Flow cytometry gating scheme. Viable macrophages are gating as CD45+GhostDye-CD11b+F4/80+

**Sup Fig 3.**
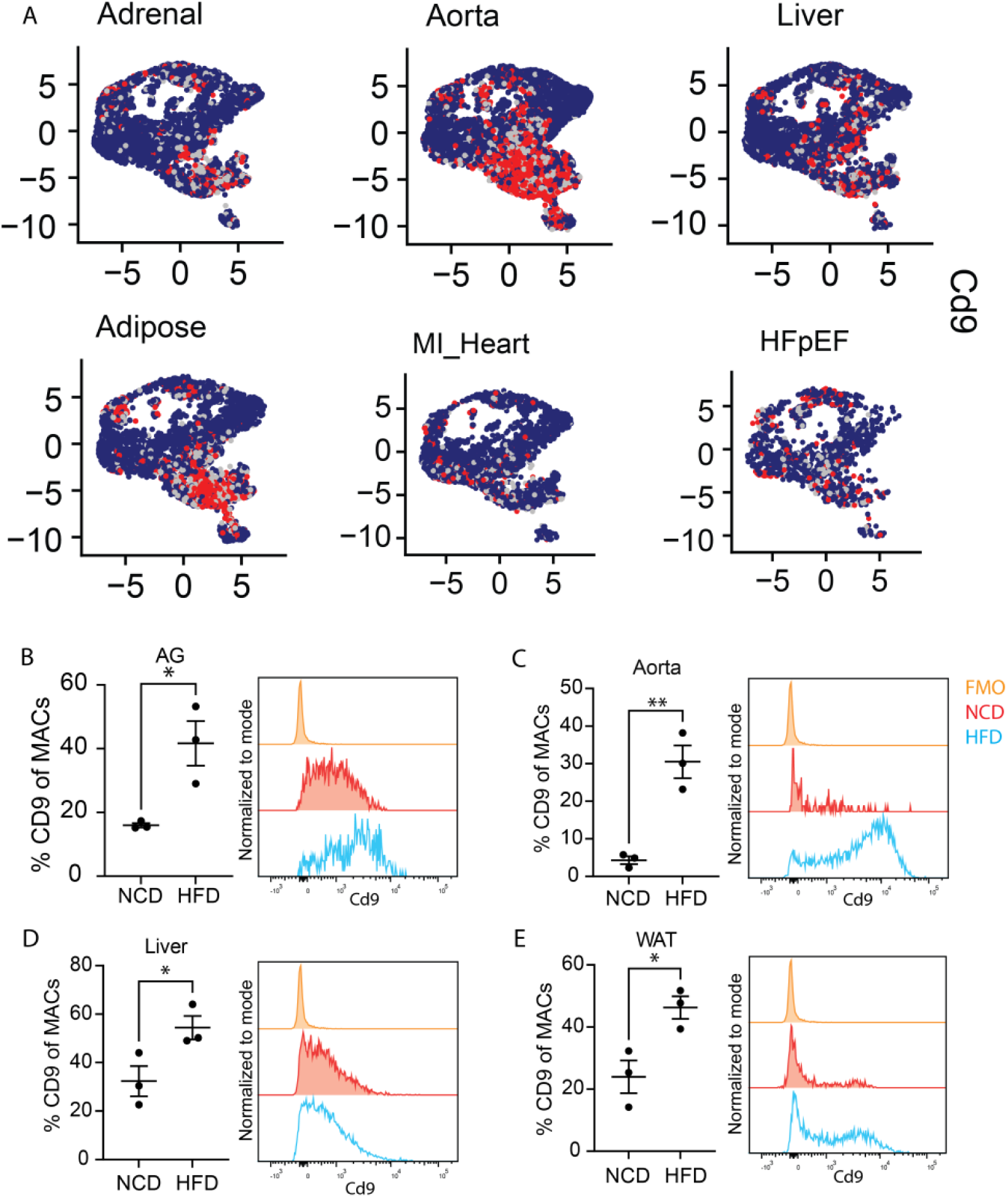
Validation of LAM features using flow cytometryA. Cd9 expression in mouse tissues. Quantification and histogram of CD9+ macrophage in adrenal gland (**B**), aorta (**C**), liver (**D**) and white adipose tissue (**E**).

